# A method for morphological feature extraction based on variational auto-encoder : an application to mandible shape

**DOI:** 10.1101/2022.05.18.492406

**Authors:** Masato Tsutsumi, Nen Saito, Daisuke Koyabu, Chikara Furusawa

## Abstract

Shape analysis of biological data is crucial for investigating the morphological variations during development or evolution. However, conventional approaches for quantifying shapes are difficult as exemplified by the ambiguity in the landmark-based method in which anatomically prominent “landmarks” are manually annotated. In this study, a morphological regulated variational autoencoder (Morpho-VAE) is proposed that conducts image-based shape analysis using imaging processing through a deep-learning framework, thereby removing the need for defining landmarks. The proposed architecture comprises a VAE combined with a classifier module. This integration of unsupervised and supervised learning models (i.e., VAE and classifier modules) is designed to reduce dimensionality by focusing on the morphological features in which the differences between data with different labels are best distinguished. The proposed method is applied to the image dataset of the primate mandible to extract morphological features, which allow us to distinguish different families in a low dimensional latent space. Furthermore, the visualization analysis of decision-making of Morpho-VAE clarifies the area of the mandibular joint that is important for family-level classification. The generative nature of the proposed model is also demonstrated to complement a missing image segment based on the remaining structure. Therefore, the proposed method, which flexibly performs landmark-free feature extraction from complete and incomplete image data is a promising tool for analyzing morphological datasets in biology.

**AUTHOR SUMMARY:** Shape is the most intuitive visual characteristic; however, shape is generally difficult to measure using a small number of variables. Specifically, for biological data, shape is sometimes highly diverse as it has been acquired through a long evolutionary process, adaptation to environmental factors, etc., which limits the straightforward approach to shape measurement. Therefore, a systematic method for quantifying such a variety of shapes using a low-dimensional quantity is needed. To this end, we propose a novel method that extracts low-dimensional features to describe shapes from image data using machine learning. The proposed method is applied to the primate mandible image data to extract morphological features that reflect the characteristics of the groups to which the organisms belong and then those features are visualized. This method also reconstructs a missing image segment from an incomplete image based on the remaining structure. To summarize, this method is applicable to the shape analysis of various organisms and is a useful tool for analyzing a wide variety of image data, even those with a missing segment.

## INTRODUCTION

Morphology is one of the most intuitively recognizable phenotypes for all organisms and has been thought to result from a variety of complex factors, including adaptation to environmental factors and speciation through neutral evolution. Therefore, comparing morphology among species and individuals is expected to provide insight into the functional role of shape and its developmental and evolutionary history. To decipher such factors from the morphology, quantification and characterization of shape are critical as it allows us to describe, interpret, and visualize the variations in shape. So far, a great deal of effort has been made towards shape analysis, and various methods have been proposed. The most widely used shape analysis is landmark-based geometric morphometrics in which landmarks are defined by anatomically homologous points on multiple samples, and the shape of a given sample is characterized by the coordinates of these landmarks [1–5]. The applications of this landmark-based method are wide-ranging, including vertebrates [6–12], arthropods [13–16], mollusks [17–19], plants [20, 21]; however, there are several difficulties and ambiguities intrinsic to this method despite its prevalence. First, the landmark-based method is unsuitable for comparisons between species and developmental stages in which biologically homologous landmarks cannot be defined. Second, both a large and small number of landmarks can cause the loss of information about the morphology of a sample [3, 5, 22–24]. In addition, errors can be problematic, such as those from measurement devices [25] and setting configurations of landmarks set inadequately by researchers due to differences in skill levels [26]. Besides the landmark-based method, elliptic Fourier analysis (EFA) has also been proposed [27, 28] and applied to characterize the shape of cells [29, 30], bivalves [31], fish [7, 32, 33], and plant organs [34–36].

Typically, the landmark-based method or EFA is combined with principal component analysis (PCA) to reduce high-dimensionality in morphological data into easily visualizable low-dimensional space [1, 6, 7]. Linear methods that reduce dimensionality, such as PCA and linear discriminant analysis (LDA), are straightforward and easily implementable, but a nonlinear approach, such as deep neural network (DNN), might be suitable for capturing more complex features with fewer dimensions. In fact, nonlinear methods based on DNN have been the standard analysis tools in the fields of image classification [37, 38] and medical diagnostic imaging [39, 40]; however, their application to morphological analysis, specifically to feature extraction of morphology, has been still limited to a few cases [41, 42]. A possible drawback of the DNN approach is that the analysis is often black-boxed and difficult to interpret, but many attempts have been made to solve this issue [43–45].

In this paper, a novel and landmark-free method based on a variational autoencoder (VAE) is proposed that analyzes shape from image data without manual landmark annotation. A VAE is a class of DNN and consists of the encoder and decoder in which the encoder embeds high-dimensional image data into low-dimensional latent variables, and the decoder reconstructs the input image from the compressed latent variables [46]. The nonlinear data compressibility of the encoder has allowed VAE to be used for feature extraction from image data [47–49]. The reconstruction capability of decoder also ensures that the input image is compressed while maintaining the information of input image, rather than being compressed in an irreversible way. Herein, the original VAE is modified by integrating a classifier module into the VAE, which allow us to extract morphological features that can best distinguish data with different labeled classes.

The modified VAE model is demonstrated to be superior than the original VAE and PCA-based methods in capturing morphological features by analyzing the mandibular image data of primates. The mandible varies widely in morphology depending on its function and diet [50–53]. For instance, the size and morphology of the mandible joint and its position relative to the biting surface differ between carnivorous and herbivorous mammals due to the differences in their masticatory functions [54, 55]. The proposed method provides a landmark-free and non-linear feature extraction analysis for the morphological data of a 3D object, as exemplified by the mandible. Additionally, an interpretation of the extracted features is presented as well as the application to the mandibular image data with a missing bone segment. The proposed model is a useful and flexible tool for investigating a morphological dataset.

## RESULTS

The study aims to develop a landmark-free method for extracting morphological features from images to distinguish different groups. A total 147 mandibles samples from seven different families (i.e., 7 labels) are prepared for verifying the method. These samples comprise 141 samples of the primate mandibles (Cercopethecidae, Cebidae, Lemuridae, Atelidae, Hylobatidae, and Hominidae) and six samples of the mandibles of carnivora (Phocidae) as an outgroup. The corresponding 3D mandible data are projected from three directions to produce three projected 2D images, as shown in Fig 1A (see Method section). These three projections of each mandible are used as the input images for the following analysis. The proposed architecture, morphological regulated variational auto encoder (Morpho-VAE), is illustrated in Fig 1B. Note that the VAE module is combined with the classifier module through the latent variable *ζ*. Subsequently, learning is performed based on the loss function *E_total_* = (1 – *α*)(*E_VAE_*) + *αE_C_*, where *E_VAE_* is the loss associated with VAE (i.e., the reconstruction + regularization losses), *E_C_* is the classification loss for the classifier module, and *α* is a hyperparameter that dictates the ratio between *E_VAE_* and *E_C_* in *E_total_*. Using the mandible sample images, the hyperparameter *α* is determined as 0.1 through cross-validation (Fig 1C, see also Method section). This choice of *α* ensures a low *E_C_* with a negligible increase in *E_VAE_* from *α* = 0, indicating that the classification ability can be incorporated into the VAE without lowering the performance in the VAE module. Other hyperparameters, such as the number of layers, number of filters, type of activation function, and optimization function, are also tuned; moreover, the number of dimensions of the latent variable are set to three (see Methods section).

**Fig 1.**
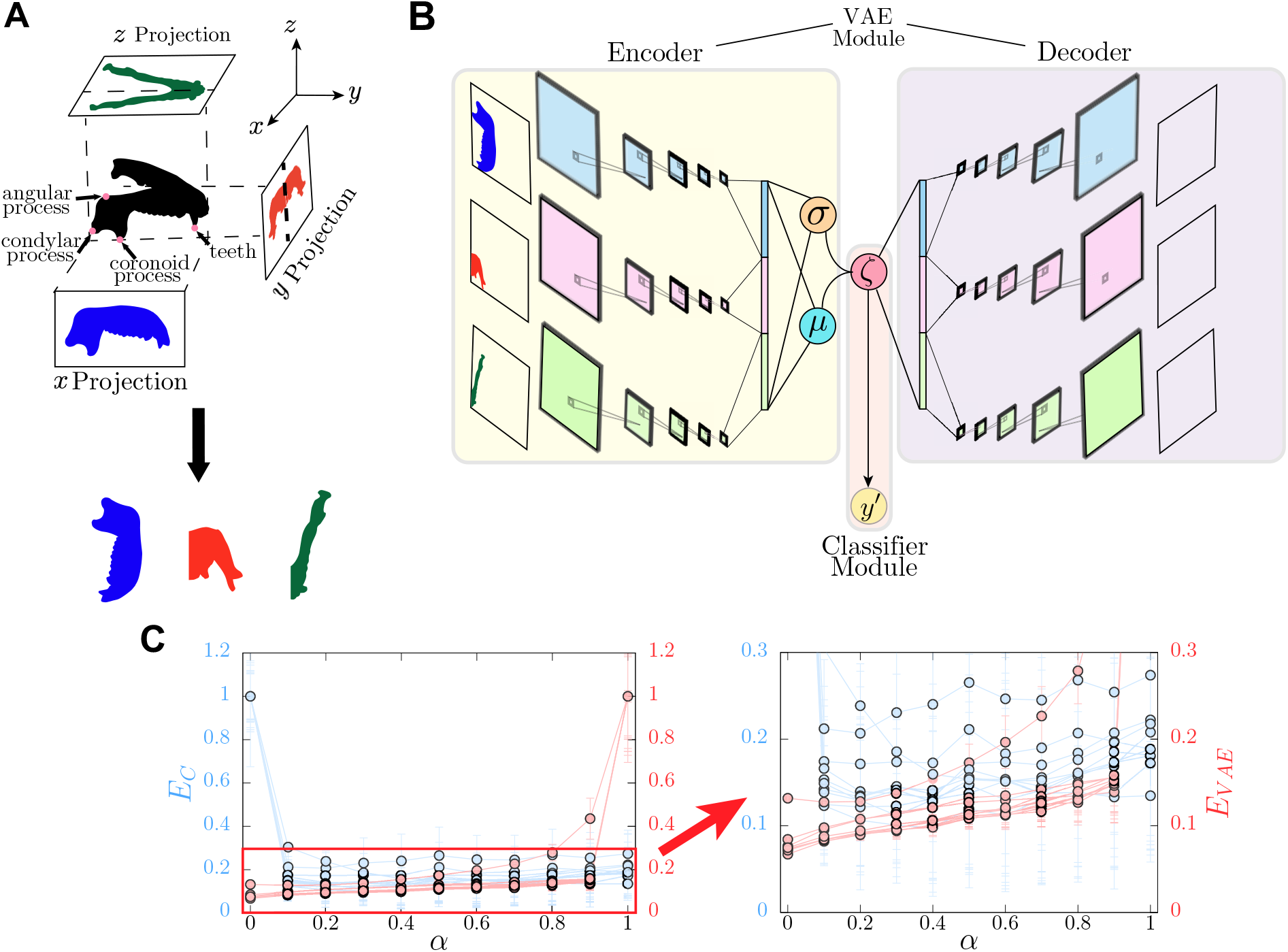
Machine learning pipeline for predicting. (A) Schematic of data preprocessing. (B) Schematic of the Morpho-VAE that comprises the encoder, decoder, and classifier. (C) Plot showing the changes in *E_C_* and *E_VAE_* as α is varied: Blue points and red points indicate the values of *E_C_* and *E_VAE_*, respectively, in the optimal model for each of the 10 combinations of training and test data. *E_C_* and *E_VAE_* are normalized such that the maximum value is 1. The left panel shows the range from 0 to 1, and the right panel shows the expanded range from 0 to 0.3.

### Cluster Separation

After the 100-epoch training as described in the Method section, a trained model is obtained that can classify the input image into seven class labels with a high validation accuracy (90% as median, SFig 2B), compress the image into 3D latent space ζ, and reconstruct the image from the latent space. The distribution of training and validation datasets in the latent space (Fig 2A) illustrates that the data points of each label form well-separated clusters from the data with different labels, compared to the results from the dimensionality reduction performed using PCA (Fig 2B) and VAE (Fig 2C). Herein, PCA is performed by transforming the image into a vector of 16,384 (= 128 × 128) dimensions and extracting the top three components; VAE is trained using the same procedure and training, validation and testing datasets to Morpho-VAE as described in the Methods section while ignoring classification loss (i.e., *α* = 0). To quantify the extent to which the data points with different class labels are separated in each method, the cluster separation index (CSI) is defined as follows:

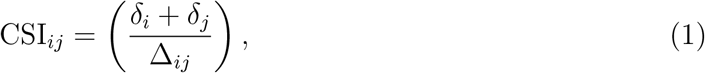

where 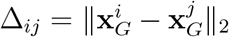 is the Euclidean distance between the centroids of the *i*-th cluster *C_i_*, 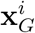, and the *j*-th cluster *C_j_*, 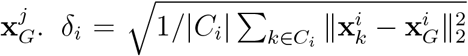 is the mean distance between a point in *C_i_*, 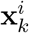, and the *i*-th cluster centroid, 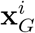. When the clusters *i* and *j* are separated, CSI_*ij*_ < 1, and CSI_*ij*_ > 1 when one of the clusters is encompassed or partially overlaps the other one. By taking the average of the maximum of CSI_*ij*_ for *j* ≠ *i* (i.e., 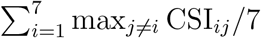), this index corresponds to the Davies-Bouldin index with *p* = *q* = 2 [56], which is widely used to evaluate the degree of cluster separation. Fig 2D shows the CSIs for all pairs of the seven clusters obtained in the reduced feature space of Morpho-VAE, PCA, and VAE, in which a single circle indicates a pair of different classes. Consistent with this result, all points are less than one, which indicates that all pairs of clusters are well-separated; however, for PCA and VAE, almost half of all points are lower than one, suggesting that the data points with different family labels cannot be distinguished in PCA or VAE space. For further verification, the evaluated Davies-Bouldin indices (a score of less than one represents well-separated clusters) are 0.74 (Morpho-VAE), 1.85 (PCA), and 2.09 (VAE).

**Fig 2.**
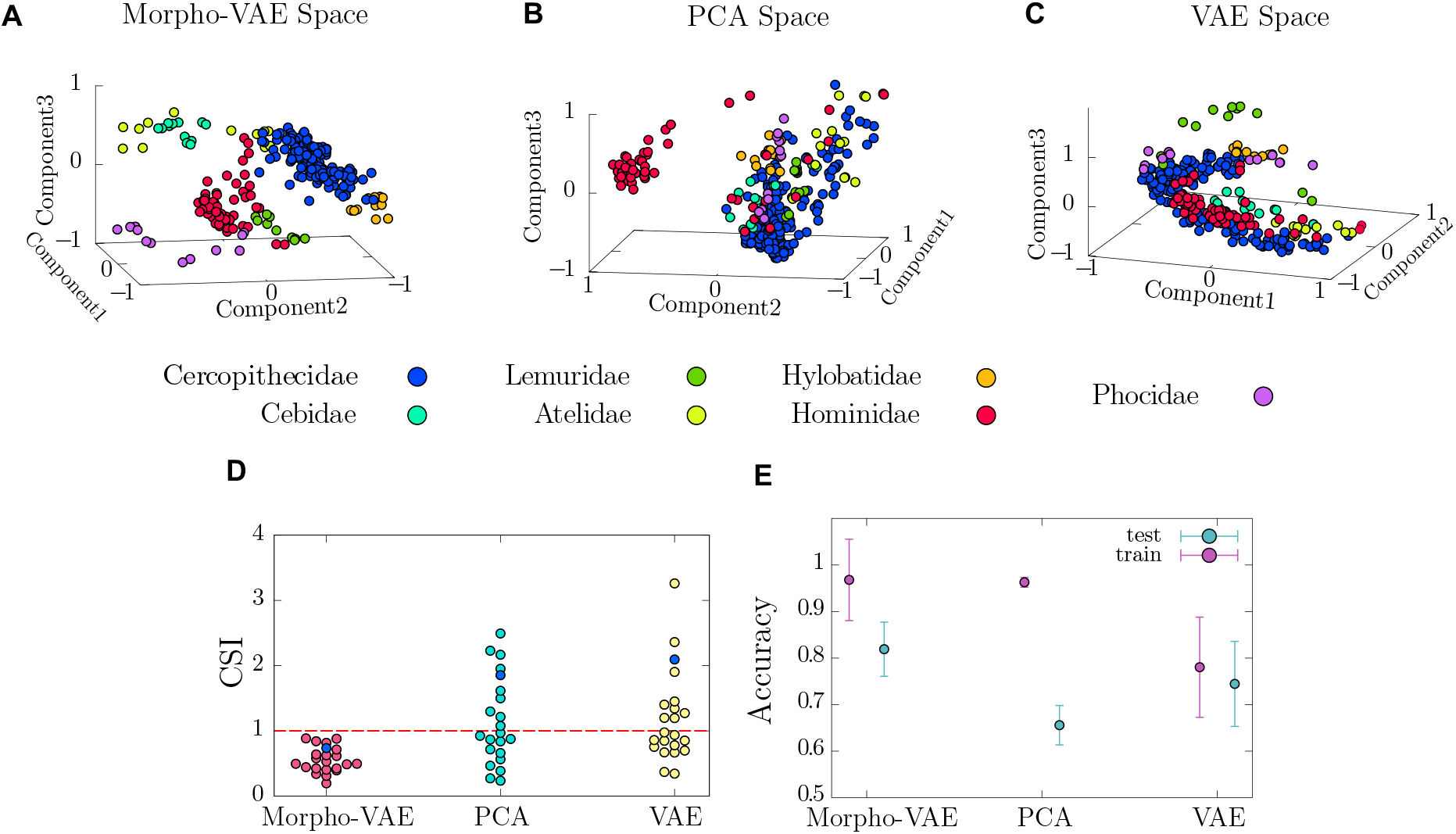
Distribution of data in latent space. (A-C) Data distribution in latent space: By using the Morpho-VAE, PCA, and VAE, respectively, all the input images are the same and are dimensionally compressed into a 3D latent space by each of the methods. (D) Dot plot of CSI a point below 1 represents a pair of well separated clusters: Blue dots represent Davies–Bouldin indices for different models. (E) Classification accuracy of families by SVM as a measure of cluster separation: The error bars indicate the mean and standard deviations in the accuracy for each of the 10 tuned models.

Additionally, the classification accuracy calculated by the support vector machine (SVM) from the data distribution in the latent space is quantified as another measure of the degree of cluster separation. Since the SVM can solve a classification problem with a high validation accuracy when the clusters of data with different labels are well-separated in the latent space, this SVM-based accuracy is expected to reflect the degree of cluster separation. After the proposed Morpho-VAE is trained using the training data (for PCA, the top three PC vectors from the training data are selected), the same training data is used for training the SVM, and then the SVM accuracy in the latent space is calculated using the test data. The average test accuracy estimated from 10 different combinations of training and test data is shown in Fig 2E in which the Morpho-VAE model achieves a much higher test accuracy than other methods, indicating that the proposed model can embed the data of different families in well-separated clusters in latent space.

### Reconstructing and Generating Images from Latent Space

The proposed Morpho-VAE model can reconstruct an image from the low-dimensional latent variable *ζ* through the decoder as well as compress the input image into *ζ* through the encoder. This ability guarantees that the compressed latent variable *ζ* preserves the information about the morphology of the input data, rather than compressing it in an irreversible manner. A representative example of an input and reconstructed images from the input image is shown in Fig 3A in which the entire morphological information of the input image is preserved in the reconstructed image, and some detailed differences are recognizable. The reconstruction loss *E_Rec_* that reflects the accuracy of the reconstructed input image reaches a plateau during training (SFig 2C), indicating that learning is successful. The reconstructed image is re-inputted into Morpho-VAE to further confirm the extent of morphological information preserved in the reconstruction image; subsequently, the predicted label is obtained through the classifier module and the prediction accuracy is calculated by comparing with the true label. This prediction accuracy can be used as an indicator of the extent of morphological information that is preserved as the precisely reconstructed images should be correctly classified, but the poorly reconstructed images should result in a significant accuracy drop. Fig 3B illustrates this prediction accuracy of the reconstructed image in comparison to the accuracy calculated from the original data with only a few percent of drops observed. Therefore, the reconstruction is demonstrably successful.

**Fig 3.**
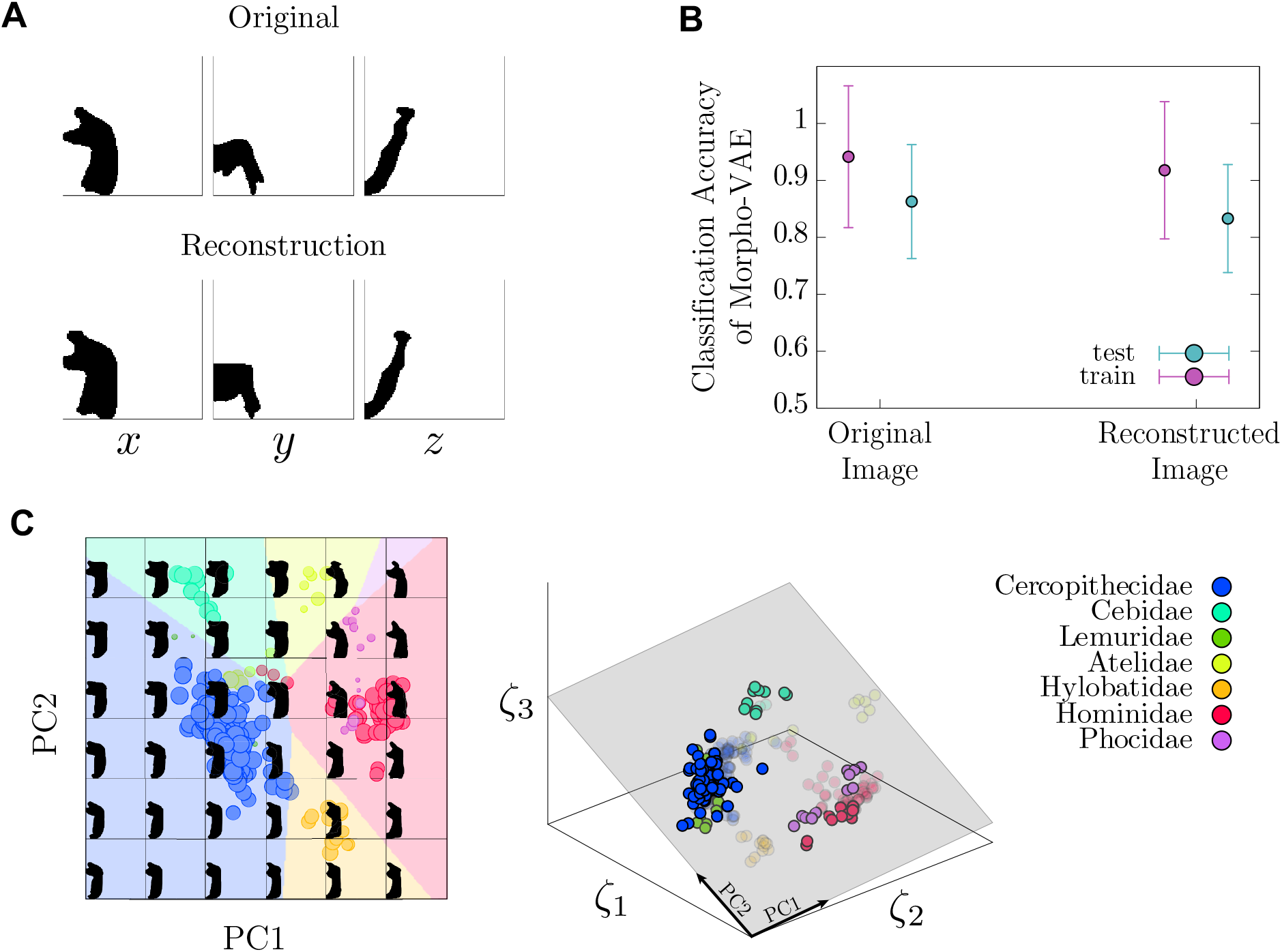
Image reconstruction by the proposed Morpho-VAE. (A) Comparison between the original and reconstructed image. (B) Classification accuracy of reconstructed images: The error bars indicate the mean and standard deviations in the accuracy for each of the 10 tuned models. (C) Generating images from latent space: Left figure shows the images reconstructed from the grid points in the PC plane at PC3 = 0 in the Morpho-VAE space, and each point is a data point projected from the Morpho-VAE space onto the PC plane. The size of each point is proportional to the absolute value of its distance from the PC3 = 0 plane. The larger the size, the closer the point is to the PC3 = 0 plane. Right figure shows the positions of points with regards to the PC plane (gray colored).

Similar to VAE, the Morpho-VAE model is categorized as a class of generative models that can generate an image from an arbitrary point in the latent space *ζ* even when no input data corresponds to the point in *ζ*. This property enables visualization of the latent space; Fig 3C illustrates the generated images from the uniformly sampled *ζ* on the 2D square lattice in 3D latent space (right panel in Fig 3C) in which the choice of the 2D plane in the 3D latent space is determined by PCA based on the data distribution in the latent space. The background colors in the left panel of Fig 3C represent the predicted labels from *ζ* by the classifier module; circles indicate the input data points mapped into *ζ* with their sizes corresponding to the distance from the PC1–PC2 plane. The generated morphology changes gradually in the latent space (left panel in Fig 3C), indicating that a smooth embedding is achieved of the morphological information into the latent space. In addition, both PC1 and PC2 seem to reflect an anatomical meaningful feature since the angle between the condylar and the coronoid processes approaches 90 degrees as PC1 becomes larger (left panel of Fig 3C), and the angular process becomes larger as PC2 increases.

### Visual Explanation of the Basis for Class Decisions

An interpretation of which part of the image Morpho-VAE focuses on the classification task can be made. Herein, a post hoc visual explanation method Score-CAM [45] is used for visualizing important areas in the input image for classification. The schematic overview of Score-CAM is given in SFig 3 (see Method section for detailed procedures). Outcomes of this analysis are “the saliency maps” for each family as shown in Fig 4A in which the darker colors represent the area judged more important for classification by the Morpho-VAE. These maps emphasize essential bone processes: the area around the coronoid process (Fig 1A) for Phocidae, the condylar process for Cercopethecidae, Hylobatidae, and Hominidae. Futhermore, the angular processes, except for Hylobatidae, are highlighted in the *x* and *y* projections. These processes connect temporal and pterygoid muscles as well as are crucial in the opening and closing of the jaw; therefore, them being highlighted for classification is reasonable.

**Fig 4.**
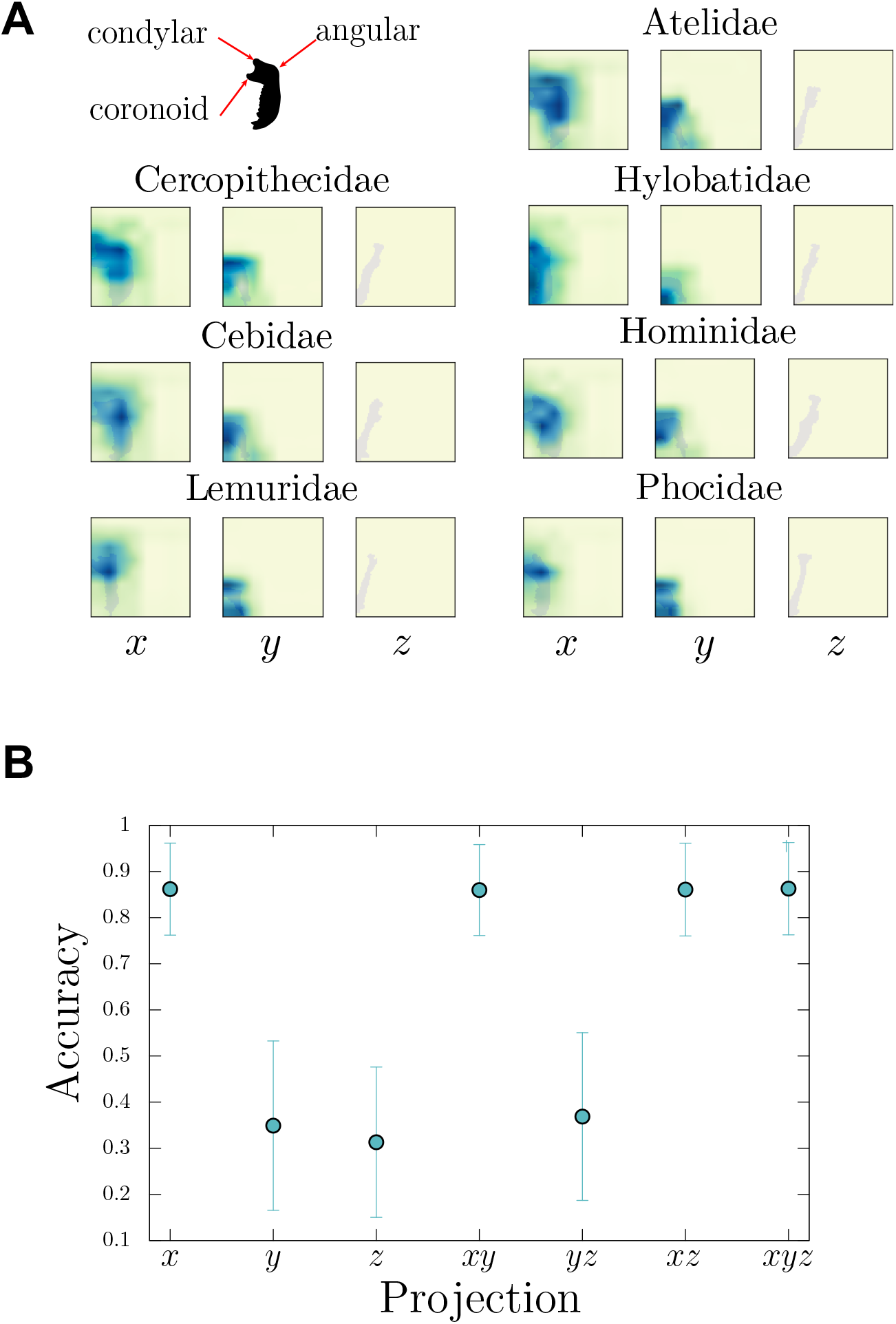
Visualization of the saliency map by Score-CAM. (A) Saliency map in each family calculated by the Score-CAM method: The stronger the color, the more intensively the area is highlighted for classification. (B) The horizontal axis is the projection direction used for the input image (e.g., *xy* indicates that the input image in the *z* direction is a blank image). The vertical axis refers to the class classification accuracy with the input. The error bars indicate the mean and standard deviations in the accuracy for each of the 10 tuned models.

The Score-CAM analysis also clarifies that the images of *z* projection do not contribute to the classification task as the colormaps in z projection are all blank (Fig 4A). This result is further confirmed by calculating the classification accuracy from the inputs of single-direction data only (e.g., *x* projection only) and those of double-direction data only (e.g., *x* and *y* projections only), rather than the full dataset of *x*, *y*, and *z* projections (Fig 4B). Both results indicate that the *x* projection image is most informative. Likewise, the site around the teeth in the *x* projection (bottom half of the image) tends to be ignored by the map, which likely reflects that the position of the teeth and their presence/absence varies greatly among samples and is thus less informative.

### Reconstruction from Cropped Data

Bone samples, especially fossil samples, sometimes miss a part. A possible application of the generative ability of the proposed model is to reconstruct such missing bone parts based on the remaining parts. Herein, the proposed model is demonstrated to achieve this reconstruction from a partially cropped image. Artificially cropped 3D data from the *y* and *z* directions (Figs 5G and 5J) is prepared and their *x*, *y*, and *z* projections are used as the data set to be reconstructed. Figs 5A, 5C, and 5D show representative examples of the original, vertically cropped, and horizontally cropped data, respectively, and their reconstructions by the proposed Morpho-VAE are presented in Figs 5E (vertical crop) and 5F (horizontal crop). The reconstructed images from the cropped data (Figs 5C and 5D) illustrate that the cropped area in the mandible of the original image (Fig 5A) is reconstructed well but not perfectly. The image looks closely similar to the reconstructed image from the original (Fig 5B), indicating that the cropped region is less informative than the remaining region.

**Fig 5.**
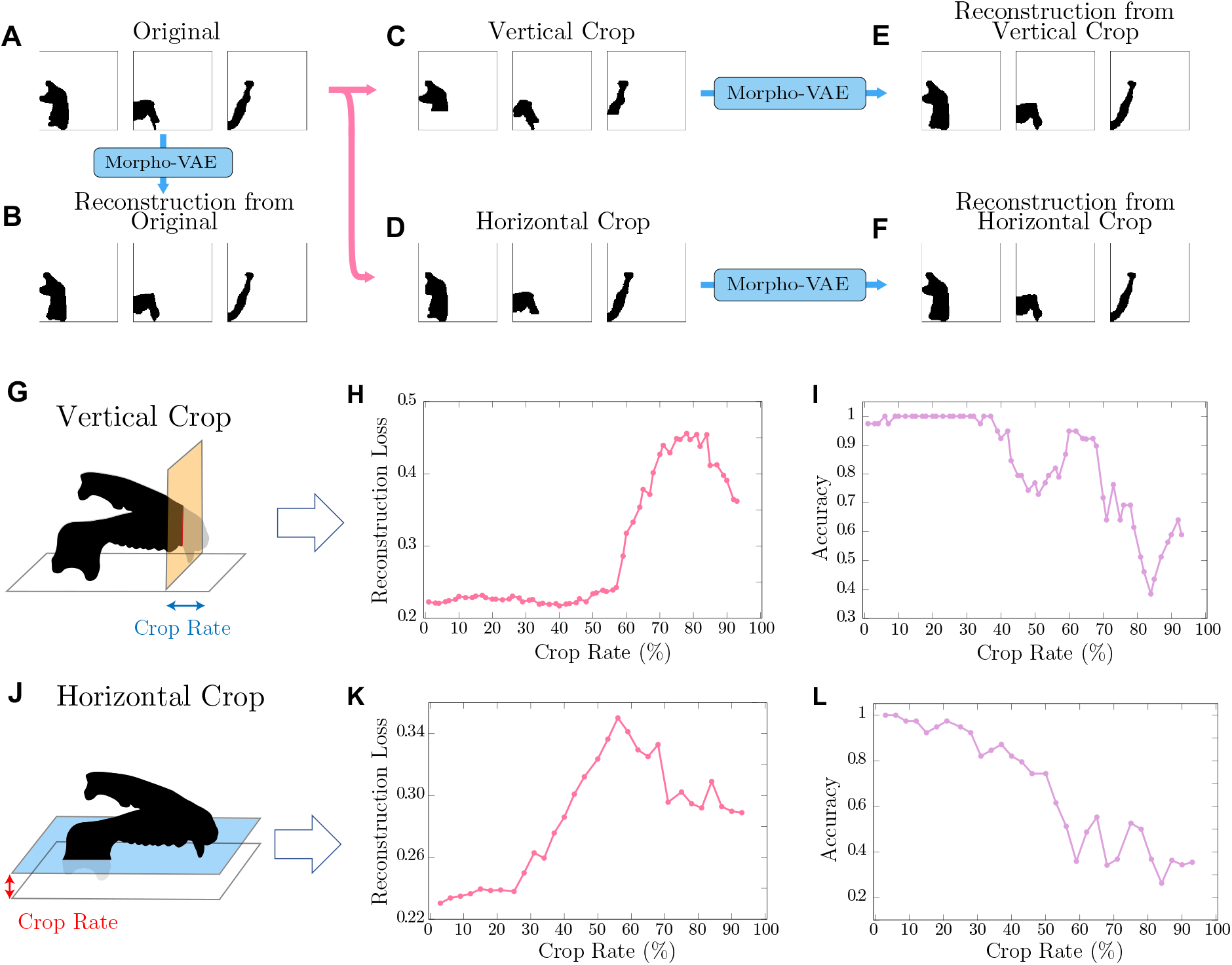
Reconstruction of cropped image. (A–F) Procedure of image cropping and reconstruction: The figures are the mandibles of *Homo Sapience*. The panels (C) and (D) represent 40% vertical and 32% horizontal cropping, respectively. (G–L) Reconstruction loss and accuracy after vertical and horizontal image cropping: Crop rate is the percentage of the mandible missing relative to its vertical or horizontal length. The figures in the first column exemplify the cropping of the mandible data. The graph in the second column shows the reconstruction loss between the reconstructed and original images. The graph in the third column shows the classification accuracy of the reconstructed images.

Furthermore, the robustness of this reconstruction is evaluated by calculating the cropped region dependency of the reconstruction loss, i.e., the binary cross-entropy between the reconstructed image from the cropped data and the original image (Figs 5H and 5K, respectively) as well as that of the prediction accuracy (Figs 5I and 5L). Within about 60% and 25% crop rates for the vertical (Figs 5H and 5I) and horizontal (Figs 5K and 5L) crops, respectively, only a slight increase in the loss and drop in the accuracy is observed, indicating that the reconstruction quality is maintained. The loss then starts to increase and the accuracy drops for further increases in the crop size. For the vertical crop, an image with the cropping size just before the loss starts to increase is shown in Fig 5D in which the shapes of the coronoid and condylar processes are just barely preserved. When these processes are completely removed, the reconstruction and classification fail (SFig 4). For the horizontal crop, an image just before the loss increase (Fig 5C) shows that the reconstruction is robust against the cropping of the region around the teeth and tip region of the mandible (i.e., the region around the body of the mandible). Both the aforementioned results indicate that the shape of the coronoid and condylar processes contain relevant information about the overall shape of the mandible, which is consistent with the results of the Score-CAM analysis (Fig 4A).

## DISCUSSION

In this study, a method based on VAE combined with a classifier module is proposed for morphological feature extraction and analyzing the image datasets of mandibles. The proposed method compresses the 128 × 128 pixel input image data into 3D latent space in which the data points of different families form well-separated clusters and the degree of cluster separation outperforms those obtained by VAE and PCA (Fig 2). Since the label information of image data is used as the supervisory signal for the classifier module, the proposed model incorporates the essence of supervised learning as well as that of unsupervised learning of a VAE module. This architecture is designed to reduce dimensionality by focusing on the morphological features through which the differences between predefined labels (i.e., family classes) are best distinguished. This is in contrast to VAE in which the latent variables are selected by focusing on the features common to the entire data. Consequently, the proposed Morpho-VAE can be interpreted as a nonlinear version of LDA that is designed to find a linear combination of features that separates data with different classes. While hybrid architectures of VAE-based unsupervised learning and classifier module have been investigated for solving classification tasks on a small number of labeled data with a large number of non-labeled data [57–59], the proposed method provides a novel framework in terms of dimensionality reduction and feature extraction.

The results in Fig 1C also indicate that the reconstruction loss shows negligible increase after taking into account the classification loss as depicted in Fig 1C with *α* = 0 (reconstruction only) and *α* = 0.1 (reconstruction and classification), suggesting that the reconstruction performance can be maintained to some extent by adding the classification function; moreover, this ensures the cluster separation of different label data in the latent space.

The characteristics of this model, which select the latent space that distinguishes predefined labels, can be described as extracting morphological features by focusing on synapomorphic traits through which a clade is well distinguished from others. The distance in the latent space is then considered to be a measure that contains information about these traits. Our naive expectation was that the latent space distance corresponds to the evolutionary distance. However, no clear correlation between the latent space distance and phylogenetic distance that is estimated in [60] is noted (SFig 5). This may account for the lack of datasets for families across the entire primates as well as the insufficient data within families. Even for sufficient data, the relationship between morphological and evolutionary distance is not straightforward as other factors, such as diet (carnivore or herbivore) and presence of a predator, can affect morphology more strongly than evolutionary distance, as exemplified by convergent evolution.

Furthermore, the Score-CAM method, which provides an interpretable visualization of the parts of an image that are important for classification (Fig 4), is applied to overcome the difficulty of interpreting DNN-based analysis. The first notable result of this analysis is that the *x* projection of the mandible image data is the most important for classification among the *x*, *y*, and *z* projections. This result is likely attributed to the fact that the area of the x projection is the largest and the results of Score-CAM, which focuses on the lateral view of the mandible is consistent with the previous studies in which the landmarks visible from the lateral view of the mandible are important in detecting sexual dimorphism [61–63] and inter-period variation [64]. Moreover, the analysis through a closer look at the x projection shows that the anatomically distinguishable projections of bone, i.e., the angular, condylar, and coronoid processes, are highlighted. For all groups except for Hylobatidae, the angular process is highlighted, but the condylar process for Cercopithecidae, Hylobatidae, and Hominidae are exaggerated. The angular and coronoid processes provide insertion sites for the medial pterygoid and temporalis, respectively; both of which are critical for producing bite force [55]. The coronoid process provides the temporomandibular joint, which works as the fulcrum during biting. The highlighted parts essentially correspond to key regions related to mastication; thus, them being highlighted seems reasonable. For Phocidae, the area around the coronoid process is emphasized. This is reasonable because a well-developed temporalis is a key feature of carnivora, and the coronoid process to which the temporalis inserts is notably enlarged compared with the other two processes.

As an application of the generative aspect of the model, the proposed model is demonstrated to complement a missing bone segment from an artificially cropped image (Fig 5) based on the remaining structure. The reconstruction is robust against the cropping of the region around teeth and tip of the mandible (Figs 5C, 5H, and 5I), but sensitive to the lack of the mandibular joint, i.e., the coronoid and condylar processes (Figs 5D, 5K, and 5L). Both these results are consistent with the results of the Score-CAM analysis (Fig 4A) in which the shape of the bone processes contains relevant information about the overall shape of the mandible. The proposed model can reconstruct a missing segment from data with defects, i.e., data in which a part of the sample is missing or damaged, as is often the case with fossils, as well as classify and reduce dimensionality for visualization from such datasets containing defects. This flexibility of the model is in contrast to the landmark-based method for which the application to data with defects is difficult as the landmarks of the missing parts cannot be defined. The generative model based on VAE has also been applied to jaw reconstructive surgeries for completing the missing segments of the bone based on the remaining healthy structure [65]. The proposed architecture by combining a VAE and classifier module provides a new framework for reconstructing missing bone segments while performing dimensional reduction for visualization and classification. In summary, the proposed model enables dimension reduction and feature extraction by which different label data are well-separated, providing a promising application of analyzing morphological dataset in biology. Although the model is designed for image input data, a combination with the landmark-based method is possible, for instance, the model output through Score-CAM analysis (Fig 4) can be used for defining the landmark positions in a systematic manner. The model can be extended to color image input instead of mask images, which will allow performing advanced analyses by extracting features that integrate texture information, such as bone density and fine bone depression, into shape information. The current model is based on 2D projected images of a 3D object and can be modified in the future to extend to 3D input data, which will provide a deeper analysis and higher resolution of the reconstructed image, but that will also require a huge dataset.

## METHODS

### Data Sets and Data Preprocessing

3D computed tomography (CT) scanning morphological data of primate mandibles is collected from Primate Research Institute (KUPRI) and MorphoSource.org. Phocidae is used as an outgroup, which is available from MorphoSource.org. Additionally, 3D datasets are collected, which consist of three images of the mandible captured from three orthogonal directions (i.e., top-, front-, and side-views), from Mammalian Crania Photographic Archive Second Edition (MCPA2). A total of 148 mandible datasets (87 Cercopethecidae, 6 Cebidae, 6 Lemuridae, 6 Hylobatidae, 6 Atelidae, 30 Homonidae, and 6 Phoci-dae) are collected (Table 1). Samples are restricted to full adults with no abnormalities in appearance.

**Table 1.** Family distribution

| Family | Number of Species |
| --- | --- |
| Cercopethecidae | 87 |
| Homonidae | 30 |
| Cebidae | 6 |
| Lemuridae | 6 |
| Hylobatidae | 6 |
| Phocidae | 6 |
| Atelidae | 6 |

Since typically more than 10^4^ datasets are required in machine learning for 3D images [66, 67], 2D machine learning is applied, rather than 3D, by converting 3D mandible image data into three 2D images (i.e., top-, front-, and side-views). SFig 1A shows that the mandible is aligned such that its teeth face downward, and the *xy* plane is defined as the plane to which the base of the mandible is parallel. Next, the position of the mandible is adjusted such that the line connecting the center of the two medial tips of the condylar head and the mandible tip is parallel to the *y*-axis. Since the mandibles of all animals collected in this study are left-right symmetrical, one mandible is divided into two pieces by the center of the mandible tip to increase the number of datasets moreover, one part is mirror-image inverted. The divided mandible, which is placed in the *xyz* space, is then converted into a set of three 2D images with size of 128 × 128 pixels by projection onto the *yz* (*x* projection), *xz* (*y* projection), and *xy* (*z* projection) planes. In addition, the length along the *y*-axis in the *x* projection of all the mandibles is normalized to be the same for avoiding size dependency of data (SFig 1).

### Model Description

This study aims to extract low-dimensional image features while ensuring the ability to classify the mandible images into families. To this end, Morpho-VAE (Fig 1B), a novel VAE-based model, is proposed in which a VAE module is combined with a classifier module through the latent variable *ζ*. Similar to conventional VAE, the VAE module of the Morpho-VAE model comprises an *l*-layer convolution neural network as the encoder and an *l*-layer deconvolution neural network as the decoder. The encoder is a layer for reducing the input data into low-dimensional latent variable *ζ* in which the input image is converted into the mean *μ* and variance *σ* of the multidimensional normal distribution. Subsequently, the latent variable ζ is sampled from the distribution 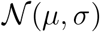. The decoder is a layer for reconstructing the low-dimensional latent variable *ζ* into an output image that has the same resolution as the input image. The network is trained such that the output image is as close as possible to the input data by optimizing the reconstruction loss *E_Rec_* (see below). The distinct feature of the Morpho-VAE is that the VAE module is combined with a classifier module in which a single layer network converts the low-dimensional latent variable *ζ* into the output vector for classification by the softmax activation function (Fig 1B). Therefore, the Morpho-VAE has two outputs: the output image for the reconstruction and the output vector for the classification. The classifier module is trained to predict the label from the input data via the latent variable ζ in a supervised learning manner. Herein, family-level classification from the input image is considered; thus, the training labels are: Cercopethecidae, Homonidae, Cebidae, Lemuridae, Hylobatidae, Phocidae, and Atelidae. A more detailed architecture of Morpho-VAE is shown in SFig 2D.

The loss functions *E_total_* that are required to train the proposed Morpho-VAE are as follows:

1. Reconstruction Loss (*E_Rec_*): binary cross entropy between the input and output images, expressed as 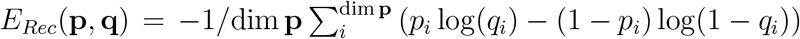, where **p** and **q** are the input and output image vectors, respectively.
2. Regularization Loss (*E_Reg_*): Kullback-Leibler divergence *D_KL_*(*q*(*ζ*|*X*)||*p*(*ζ*)) between the data distribution in the latent space *q*(*ζ*|*X*) encoded by the encoder from data *X* and the predefined reference distribution 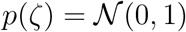, which is fixed as a Gaussian distribution with mean 0 and variance 1.
3. Classification Loss (*E_C_*): cross entropy between the predicted **y′** and true label vectors **y** from the latent variable *ζ* and classifier module, expressed as 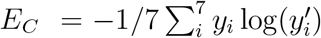.

From these three loss functions, VAE loss function is defined as *E_VAE_* = *E_Rec_* + *E_Reg_*. Moreover, the total loss function is defined as *E_total_* = (1 – *α*)*E_VAE_* + *αE_C_*, where *α* = 0.1 is selected by cross-validation (Fig 1C), and the Morpho-VAE is trained to minimize *E_total_* by backpropagation.

### Hyperparameter Tuning

The structural hyperparameters of Morpho-VAE, such as the number of layers, number of filters in each layer, type of activation function, and type of optimization function, are tuned by Optuna [68].

The number of layers is optimized to be within the range of one to five and the number of filters in each layer is optimized to be within the range of 16 to 128. The activation functions are selected from ReLU, sigmoid, and tanh, and the optimization function is selected from stochastic gradient descent, adaptive momentum estimation (Adam), and RMSprop. Note that the latent space dimension is fixed to three in these processes. These optimizations are performed by searching 500 different conditions, each with 100 epochs of training, and the following parameters are defined as the optimal hyperparameters to minimize the loss function *E_total_*. The other hyperparameters are listed in STable 2. The number of layers in the encoder is five. The numbers of filters in each layer are 128, 128, 32, 32, and 64 in the order from the layer nearest to the input layer. The selected activation and optimization functions are ReLU and RMSprop, respectively. Moreover, the number of layers in the decoder is five, and the numbers of filters in each layer are 64, 32, 32, 128, and 128 in the order from the layer nearest to the latent variable. The type of optimization function is RMSprop. Note that sigmoid is adopted instead of ReLU as the activation function of the decoder because the input image of this model is a binary image in the range of [0,1], and the output image needs to be in the same range.

After tuning the structural hyperparameters, the dimensions of the latent variable *ζ* are also explored. The number of dimensions of the latent variable is examined from 2 to 10 by 100 times independent 100-epoch training with different training-validation datasets for each dimension. SFig 2A shows that the mean and median of the minimum of *E_total_* in each 100-epoch training decrease as the dimension increases from two to eight. Since our aim is to select a low-dimensional feature ζ that generates a low *E_total_*, the dimension value of three is adopted for which only a slight increases appears in loss value compared with the dimensions ≥ 4, but a certain drop (SFigs 2A and 2B) is observed between the dimensions two and three.

A double cross-validation procedure [69] is used for separating the data into training, validation, and test data. One-third of the total data is used as test data to evaluate the generalization performance of the Morpho-VAE. Of the remaining data, 75% is separated as training data for tuning the hyperparameters of the Morpho-VAE and the remaining 25% as validation data for verifying the hyperparameters to avoid leaks of the same species of data. Since the data set collected in this study has a class imbalance as listed in Table 1, the data set is divided into training and test data using the proportional extraction method, which divides the data by reflecting the sample size of each label. Note that two datasets are obtained from one mandible sample (see Datasets section) but the data are distributed such that the same sample is not included both in the test and training data.

### Visualization of the Saliency map (Fig 4A) by Score-CAM

The Score-CAM [45] method is applied to visualize the Morpho-VAE making its decisions. The schematic overview of Score-CAM is given in SFig 3. First, upsampling is performed from the 8 × 8 pixel activation map, which activates the last layer in the convolution layers of the encoder, to a 128 × 128 pixel image and then normalization is implemented such that the maximum and minimum pixel intensities of the image are 1 and 0, respectively. Each pixel intensity of the image is then multiplied by the intensity of the corresponding pixel in the 128 × 128 original input image to create a masking image. Furthermore, this masking image is re-inputted into the Morpho-VAE and the prediction probability is calculated for the label of the input image through the classifier module. Since the calculated prediction probability can be interpreted as the importance of the masking image, this probability is then multiplied by the activation map, and the final outcome of the Score-CAM (Fig 4), “the saliency map”, is obtained by taking a sum over the number of filters (e.g., 64).

## ACKNOWLEDGMENTS

We thank: Y. Kondo, K. Aoki, Y. Himeoka, J. Iwasawa, Y. Uchida, H. Higashiyama, A. Tokuhisa, K. Terayama, and Y. Okuno for the meaningful discussions as well as M. Hasebe, T. Fujimori, T. Ueda, S. Yoshida, S. Takada, and K. Agata for supporting comments. We are grateful to T.D. Nishimura and Primate Research Institute of Kyoto University for their help in collecting CT data. This research was supported by Joint Research of the Exploratory Research Center on Life and Living Systems (ExCELLS) (ExCELLS program no. 21-319 and no. 21-102 to N.S.). This research was also supported in part by JSPS KAKENHI (17H06389 to C.F.) and JST ERATO (JPMJER1902 to C.F.). Masato Tsutsumi was supported by RIKEN Junior Research Assistant.

## SUPPLEMENTAL FIGURES

**SFig 1.**
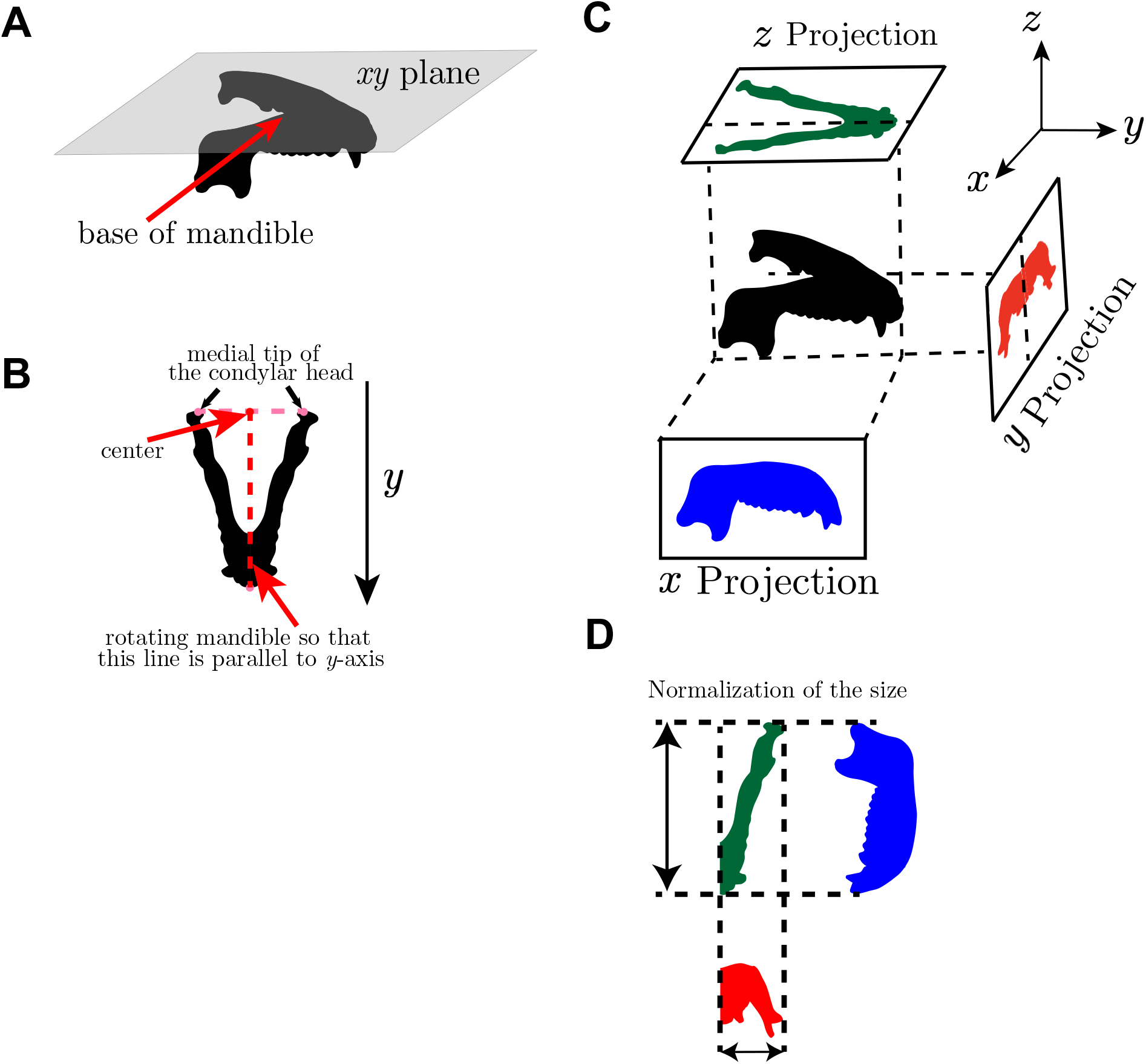
Detailed description of image preprocessing. (A) The *xy* plane is defined as a plane to which the base of the mandible is horizontal. (B) After placing the mandible in the xy plane, it is rotated so that the mandible tip and the mid-point of the line connecting the condylar head’s left and right medial tips have the same y-coordinate. (C) The mandible (arranged as shown) is projected from three orthogonal directions. (D) The size of the data is normalized such that the length from the angular process to the tip of the mandible is constant.

**SFig 2.**
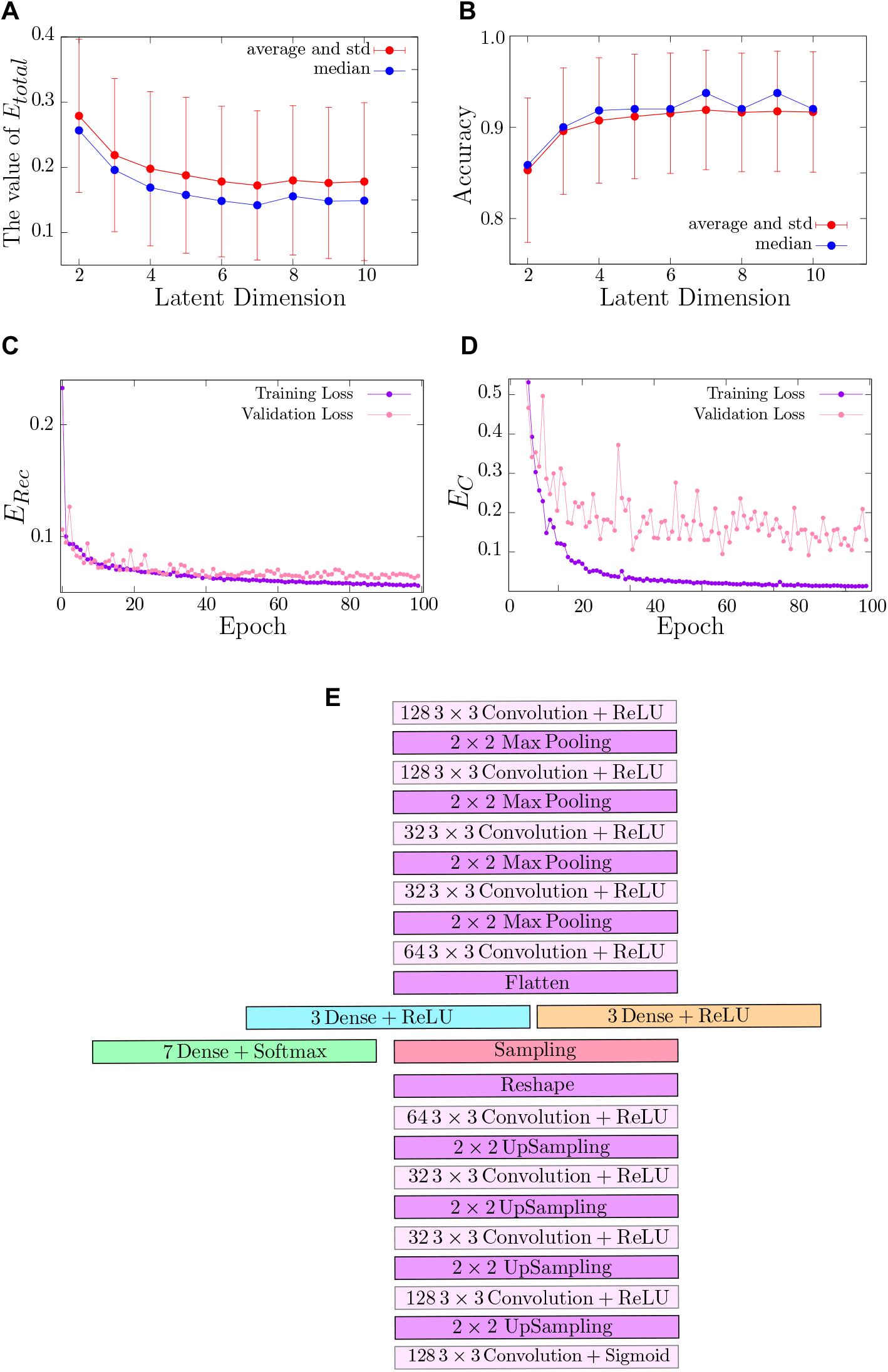
Latent dimension tuning and detailed architecture of the proposed Morpho-VAE. (A) Mean, standard deviation, and median values of *E_total_* for 2 - 10 dimension. These values are calculated from 10 independent architectures listed in STable 2. (B) Classification accuracy for 2 - 10 dimension calculated from 10 independent architectures. The selected number of dimension is 3. (C and D) Trajectories of training and validation losses for *E_Rec_* and *E_C_* during training. (E) Detailed architecture of the Morpho-VAE: The upper and lower sides of the figure are the input and output layers, respectively.

**SFig 3.**
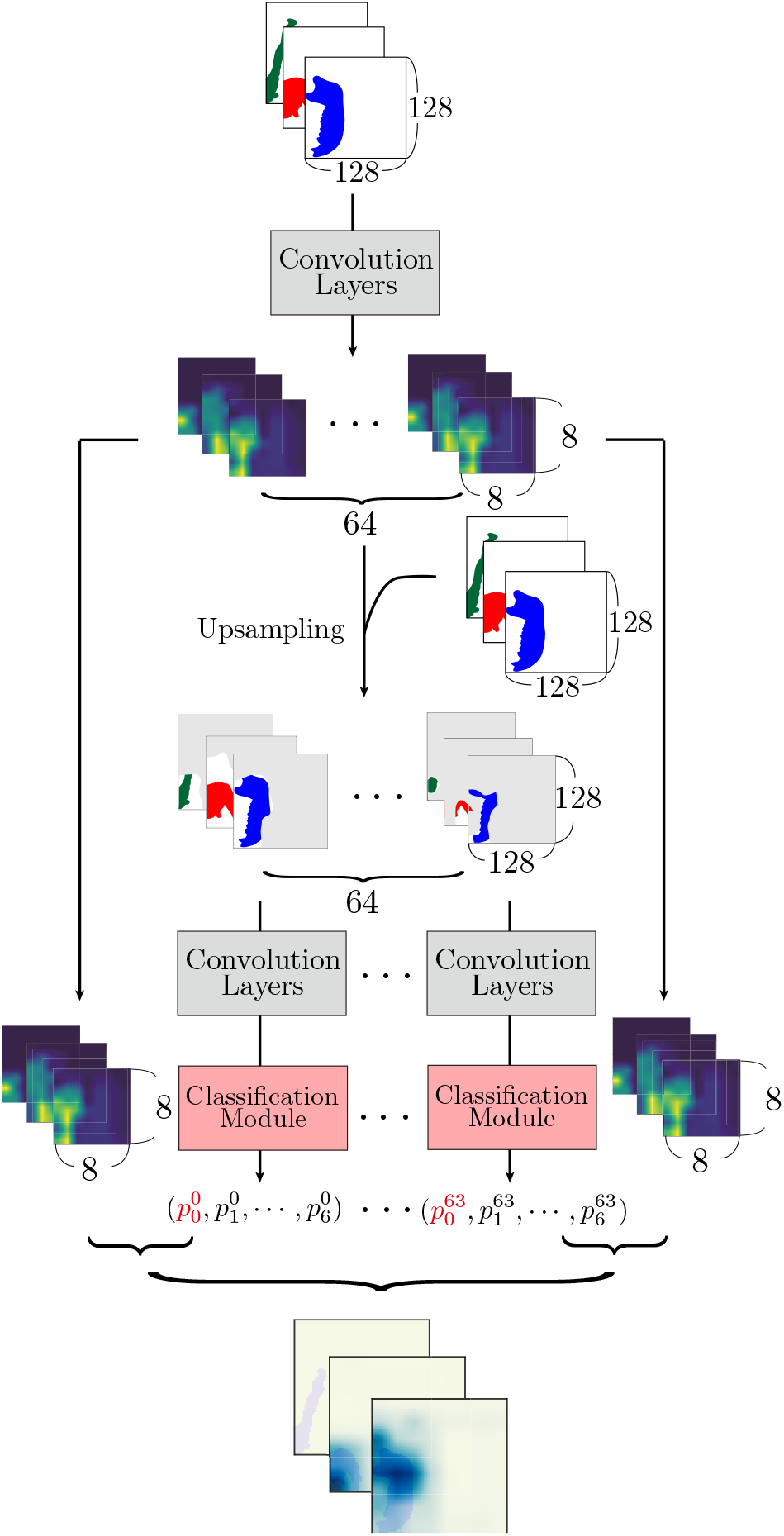
Schematic image for creating saliency map. First, the output images with 64 channels obtained from the last convolution layer of the Morpho-VAE with a set of input images are prepared. The, output images are upsampled to the same size as the input images. The resulting 64 outputs are normalized using the maximum and minimum values for each image. These 64 normalized output images are multiplied by the original input images to create 64 images. Each of these is then inputted to the Morpho-VAE to calculate the probability of the seven classes. Each of the 64 probability values is considered as the importance of the 64 outputs. Next, the importance and output are multiplied and then added together to obtain a saliency map.

**SFig 4.**
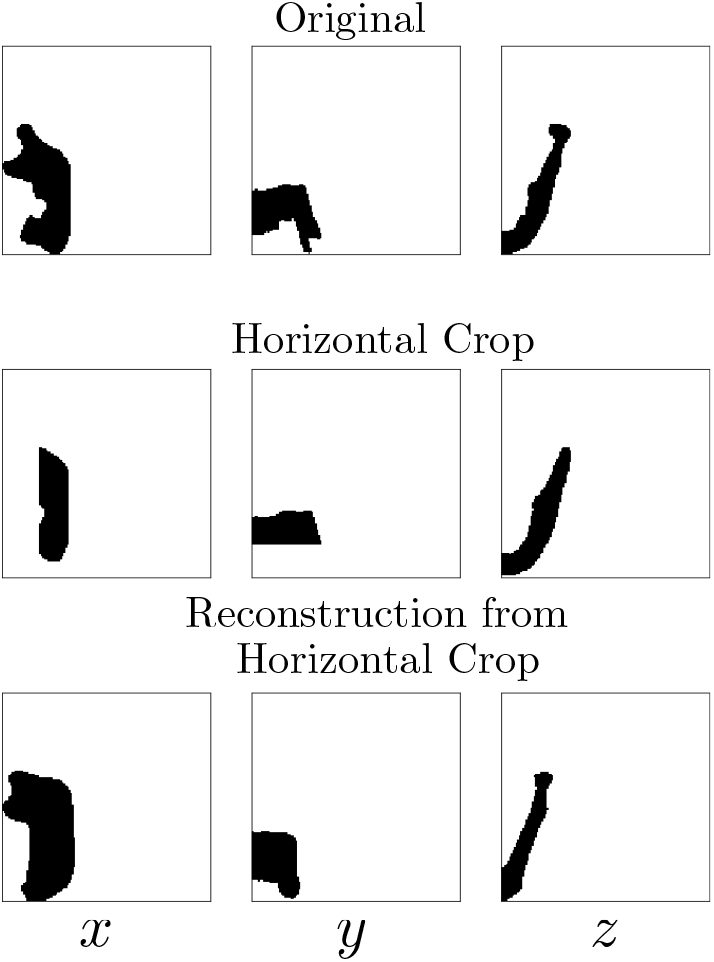
Example of reconstruction failure when crop rate is significantly large. A significant defect in the coronoid process results in the failure of the reconstruction, where renconstructed image is far different from the original image.

**SFig 5.**
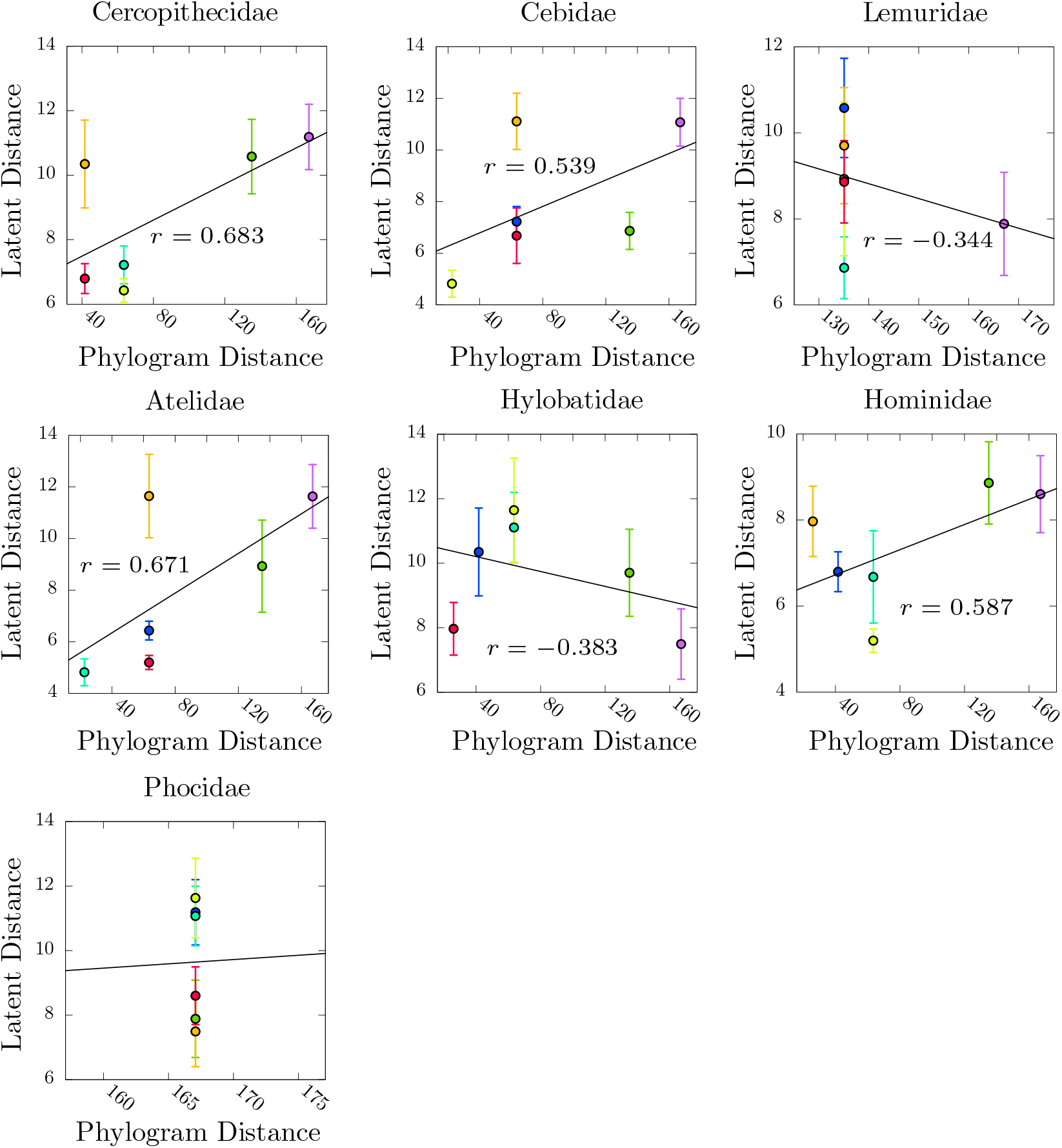
Correlation between the distance in the Morpho-VAE and phylogentic tree. The vertical axis represents the Euclidean distance between clusters in the latent space of the Morpho-VAE, and the horizontal axis represents the distance within the phylogenetic tree. The error bars indicate the mean and standard deviations in the accuracy for each of the 10 tuned models listed in STable 2.

## TABLES

**STable 1:** List of the mandible data and references.

| Family | Species | # of samples | Reference |
| --- | --- | --- | --- |
| *Cercopithecidae* | *Allenopithecus nigroviridis* | 8 | MorphoSource |
|  | *Cercocebus galeritus* | 2 | MorphoSource |
|  | *Cercopithecus ascanius* | 6 | MorphoSource |
|  | *Cercopithecus cephus* | 4 | MorphoSource |
|  | *Cercopithecus mitis* | 2 | MorphoSource |
|  | *Cercopithecus mona* | 2 | MorphoSource |
|  | *Cercopithecus neglectus* | 2 | MorphoSource |
|  | *Cercopithecus nictitans* | 2 | MorphoSource |
|  | *Cercopithecus petaurista* | 2 | KUPRI |
|  | *Chlorocebus aethiops* | 2 | MorphoSource |
|  | *Colobus angolensis* | 4 | MorphoSource |
|  | *Lophocebus albigena* | 4 | MorphoSource |
|  | *Macaca arctoides* | 4 | KUPRI |
|  | *Macaca arctoides* | 4 | MorphoSource |
|  | *Macaca assamensis* | 4 | MorphoSource |
|  | *Macaca cyclopis* | 4 | MorphoSource |
|  | *Macaca fascicularis* | 6 | MorphoSource |
|  | *Macaca fuscata* | 38 | KUPRI |
|  | *Macaca maura* | 6 | MorphoSource |
|  | *Macaca mulatta* | 2 | MorphoSource |
|  | *Macaca nemestrina* | 8 | MorphoSource |
|  | *Macaca nigra* | 2 | MorphoSource |
|  | *Macaca silenus* | 2 | KUPRI |
|  | *Macaca sylvanus* | 2 | KUPRI |
|  | *Macaca thibetana* | 2 | MorphoSource |
|  | *Macaca tonkeana* | 8 | MorphoSource |
|  | *Mandrillus leucophaeus* | 8 | MorphoSource |

STable 1: **List of the mandible data and references**
| Family | Species | # of samples | Reference |
| --- | --- | --- | --- |
|  | *Mandrillus sphinx* | 2 | MorphoSource |
|  | *Papio hamadryas* | 16 | MorphoSource |
|  | *Presbytis femoralis* | 4 | KUPRI |
|  | *Presbytis melalophos* | 2 | KUPRI |
|  | *Presbytis melalophos* | 2 | MorphoSource |
|  | *Simias concolor* | 4 | KUPRI |
|  | *Theropithecus gelada* | 4 | MorphoSource |
| *Cebidae* | *Cebus albifrons* | 2 | MCPA2 |
|  | *Cebus apella* | 4 | KUPRI |
|  | *Cebus apella* | 2 | MCPA2 |
|  | *Cebus capucinus* | 4 | MCPA2 |
| *Lemuridae* | *Eulemur mongoz* | 2 | KUPRI |
|  | *Hapalemur griseus* | 2 | MorphoSource |
|  | *Lemur catta* | 4 | MCPA2 |
|  | *Lemur catta* | 4 | MorphoSource |
| *Atelidae* | *Alouatta caraya* | 2 | MorphoSource |
|  | *Alouatta fusciceps* | 4 | MorphoSource |
|  | *Ateles geoffroyi* | 2 | MCPA2 |
|  | *Ateles paniscus* | 4 | MCPA2 |
| *Hylobatidae* | *Hylobates hoolock* | 4 | MCPA2 |
|  | *Hylobates lar* | 4 | MCPA2 |
|  | *Hylobates syndactylus* | 4 | MCPA2 |
| *Homonidae* | *Gorilla gorilla* | 8 | KUPRI |
|  | *Gorilla gorilla* | 2 | MCPA2 |
|  | *Homo sapiens* | 2 | MCPA2 |
|  | *Homo sapiens* | 40 | MorphoSource |
|  | *Pan troglodytes* | 4 | MCPA2 |
|  | *Pongo pygmaeus* | 4 | MCPA2 |

STable 1: **List of the mandible data and references**
| **Family** | **Species** | **# of samples** | **Reference** |
| --- | --- | --- | --- |
| *Phocidae* | *Zalophus californianus* | 12 | MorphoSource |

Data used in this study are listed. KUPRI, dataset from Primate Research Institute of Kyoto University; MorphoSource, dataset from Morphosource.org; MCPA2, dataset from Mammalian Crania Photographic Archive Second Edition

**STable 2:**
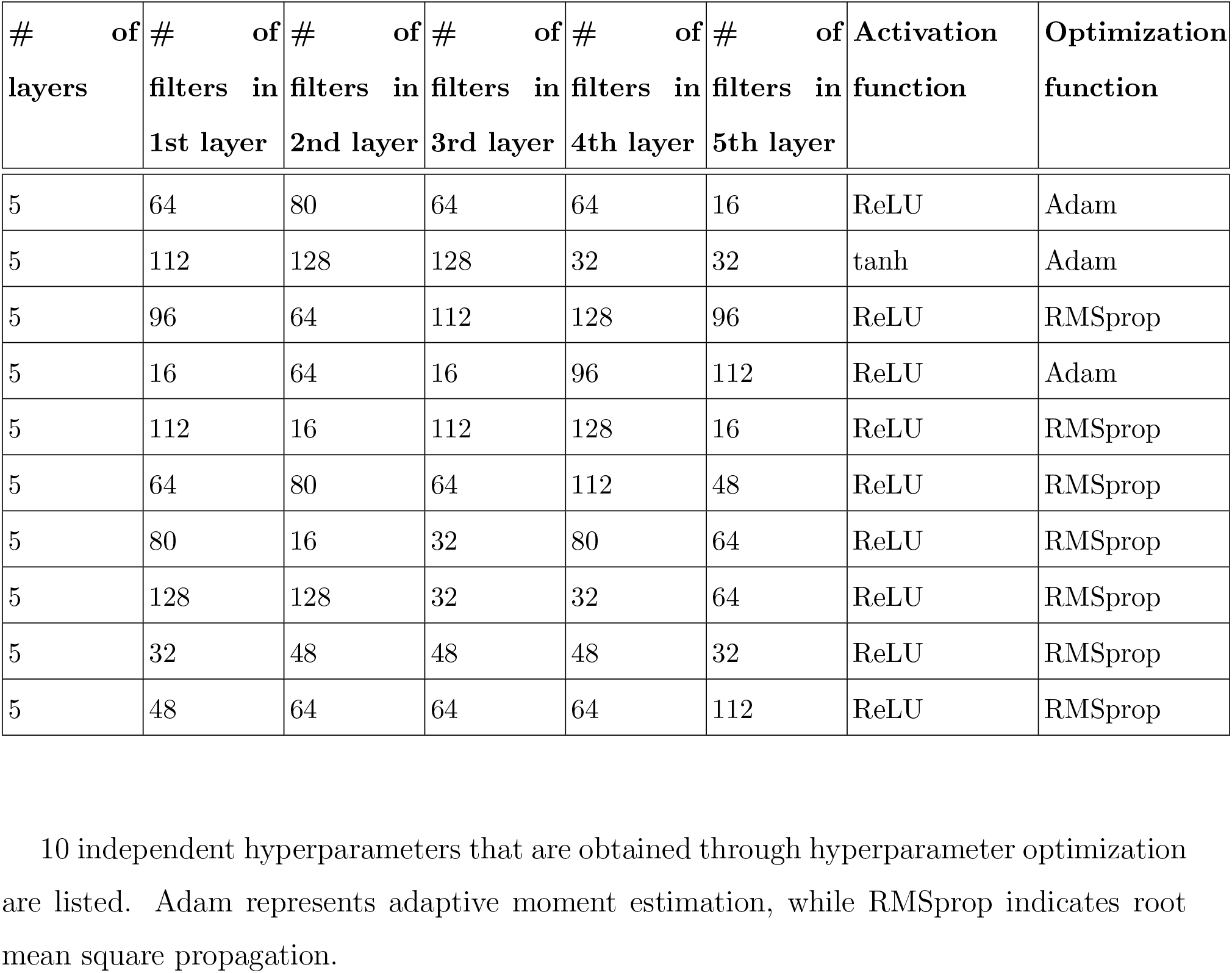
All model hyperparameters.

